## Supplementary material for "Sketch, capture and layout phylogenies": User manual

### PhyloSketch App User Manual

Daniel H. Huson
University of Tuebingen

Version 2.2.0, Nov 11, 2025

#### Contents

**1 Introduction** **2**

**2 Installation** **3**

**3 Getting Started** **3**

**4 Modes Overview** **3**

**10 Combining vs Transfer View** 13

**11 Example files** 14

**12 Advanced features** 14

**13 Support and Feedback** 14

**14 Third-Party Software** 15

### 1 Introduction

PhyloSketch App (also known as PhyloSketch2) is an application for interactively creating and editing phylogenetic trees and networks by drawing them. Written in Java using JavaFX, this program runs on macOS, Linux, and Windows and is also designed for touch-screen devices running iOS or Android.

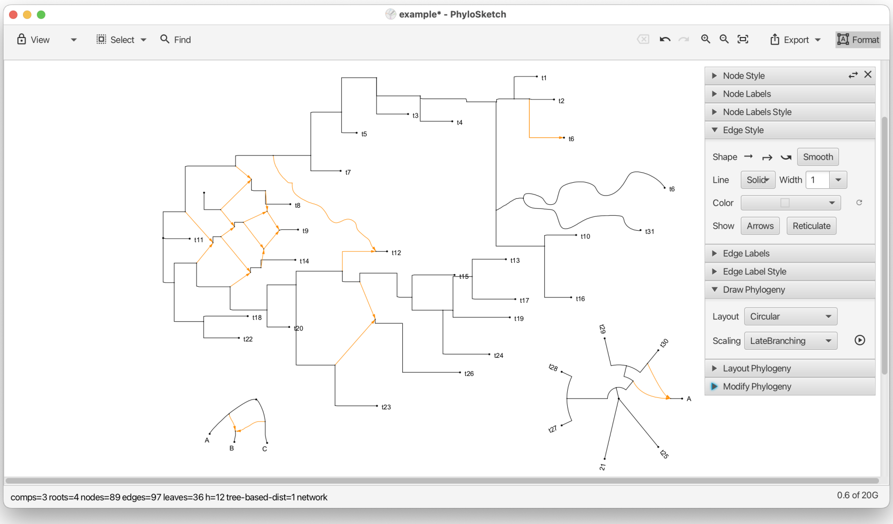

PhyloSketch App is a new program based on PhyloSketch (also known as PhyloSketch1), which was introduced in [Francis et al., 2021]. The program reuses some code from our SplitsTree app [Huson and Bryant, 2024], but most of the code and algorithms are new [Huson, 2025].

#### 2 Installation

- Desktop versions for macOS, Linux, and Windows are available from:
  - `https://github.com/husonlab/phylosketch2`
  - `https://software-ab.cs.uni-tuebingen.de/download/phylosketch2/welcome.html`
- The iOS app is available for testing via Apple TestFlight. If you are interested in beta testing, contact the author for an invitation.
- Open the app and grant any necessary permissions for accessing storage, if prompted.

#### 3 Getting Started

When you first open PhyloSketch, you are presented with a canvas containing a simple example tree. You can modify this example or start creating your own phylogenetic tree or network. The toolbar at the top provides access to all major functions, including mode selection, import options, layout, and various editing tools.

#### 4 Modes Overview

PhyloSketch operates in four primary modes. The mode can be selected using the first control on the toolbar. In the desktop version, tooltips and menu items may refer to these as *Sketch*, *Move*, *View*, and *Capture* modes, corresponding to the modes described below.

##### 4.1 Edit Mode (Sketch)

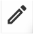

- Create new nodes and edges by drawing directly on the canvas.
- Long-press to create a new node, and press-drag to create edges.
- Shift-press-drag to move nodes or reshape edges.
- Use this mode to build a new tree or network from scratch or to modify an existing one.

##### 4.2 Transform Mode (Move)

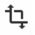

- Move nodes, reshape edges, and adjust the layout of your phylogenetic tree or network.
- Touch or click and drag nodes or edges to reposition or reshape them.

##### 4.3 Read-Only Mode (View)

- Editing is disabled, but you can still select nodes or edges to inspect labels and properties.
- Useful for viewing and presenting a tree or network without risk of accidental modifications.

##### 4.4 Capture Mode

- Use this mode when capturing a phylogeny from a background image.
- Capture nodes, edges, and labels present in the image, assisted by image-analysis and OCR.

Usually, after initial capture, Edit Mode is used to interactively complete and refine the captured tree or network.

#### 5 Tool Bar Overview

The toolbar provides access to the core functionalities of PhyloSketch:

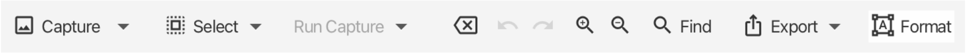

##### 5.1 Mode Selection

Select the desired mode (Edit/Sketch, Transform/Move, Read-Only/View, or Capture) using the first item on the toolbar.

##### 5.2 Selection Menu Button

Select nodes or edges based on their properties:

- **All** Select all nodes and edges.
- **None** Deselect everything.
- **Extend** Extend the current selection based on adjacency.
- **Invert** Invert the current selection.
- **Roots/Leaves** Select root or leaf nodes.
- **Articulation Nodes** Select nodes that separate biconnected components.

#### 5.3 Run Capture Menu Button

When a background image has been loaded, these items support capturing a tree or network from that image.

Pressing the Run Capture menu button once attempts to locate the root in the background image. Pressing it again performs the capture using the current parameters. Alternatively, use the following menu items explicitly:

**Load Image...** Import an image to use as a background for the network.

**Place Root** Locate the root in the background image as preparation for network capture.

**Capture Phylogeny** Extract phylogenetic structure from the image, converting it into a graph representation.

**Remove Image** Remove the currently loaded background image from the view.

##### 5.3.1 Advanced Capture Items

**Capture Labels** Use OCR to capture the labels in the background image.

**Capture Lines** Detect and capture lines from the image, typically used for network reconstruction.

**Parameters...** Open a dialog to configure various settings for network capture (line detection, OCR, thresholds, etc.).

Here is an example of a captured network [Koblmueller et al., 2007]:

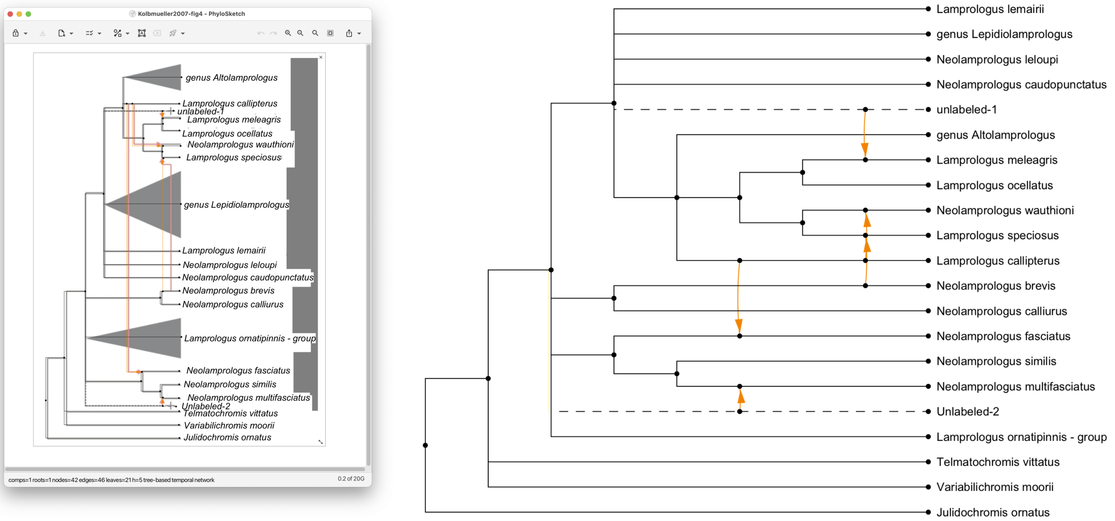

#### 5.4 Formatting Button

The formatting button toggles the visibility of the formatting pane described below.

#### 5.5 Other Toolbar Buttons

- **Delete** Remove selected nodes and edges.
- **Undo/Redo** Revert or repeat the most recent actions.
- **Zoom In/Out** Adjust the zoom level for the canvas.
- **Zoom to Fit** Zoom to fit the current phylogeny in the window.
- **Export Menu** Copy selections or save images and data in various formats.

### 6 Status Bar

The status bar reports the number of components, roots, nodes, edges, leaves, and the hybridization number $h$. It also indicates properties such as whether the current object is a *network* or a *tree-based network*.

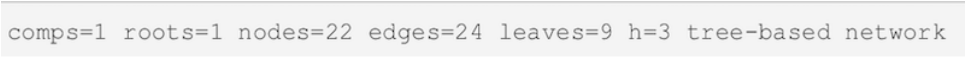

### 7 Formatting Pane

The formatting pane has several panels for formatting nodes and their labels, and edges and their labels.

#### 7.1 Node Style Panel

This panel provides options for customizing the appearance of nodes:

- **Shape** Dropdown menu to select the shape of nodes.
- **Size** Combo box to specify or edit the size of nodes.
- **Color** Color picker to select node color.
- **Revert** Button to revert a nodes style (shape, size, and color) to default values.

#### 7.2 Node Labels Panel

This panel provides options for labeling nodes:

- **Nodes to Label** Dropdown menu to select which nodes should be labeled.
- **Labeling Method** Dropdown menu to specify how the selected nodes should be labeled.
- **Unique Labels** Toggle button to enforce unique labels, preventing reuse of existing labels.
- **Label Input Field** Text field to manually enter labels for nodes.
- You can use HTML-like tags for styling text (e.g., `<i>`, `<b>`, `<sub>`, `<sup>`).

#### 7.3 Node Labels Formatting Panel

This panel provides options for formatting node labels:

- **Font** Dropdown menu to select the font for node labels.
- **Revert Font** Button to revert font settings to default.
- **Size** Combo box to specify or edit the font size for node labels.
- **Revert Size** Button to revert size settings to default.
- **Bold** (B) Toggle bold formatting for node labels.
- **Italic** (I) Toggle italic formatting for node labels.
- **Underline** (U) Toggle underlining for node labels.
- **Color** Color picker to set the text color of node labels.
- **Revert Color** Button to revert text color to default.
- **Fill** Color picker to set the background fill color of node labels.
- **Revert Fill** Button to revert the background fill color to default.

#### 7.4 Edge Style Panel

This panel provides options for customizing the appearance and properties of edges:

- **Shape** Buttons to select the shape of edges, including straight (S), rectangular (R), curved (C), and smooth styles.
- **Line** Dropdown menu to select the line style (solid, dashed, dotted, etc.).
- **Width** Combo box to specify or edit the line width.
- **Color** Color picker to set edge color.
- **Revert Color** Button to reset edge color to its default.
- **Show Arrows** Toggle to enable or disable arrowheads on edges.
- **Show Reticulate** Toggle to enable or disable orange highlighting of reticulate edges.

#### 7.5 Edge Labels Panel

This panel provides options for managing edge labels, including weights, support values, and probabilities:

- **Weight** Text field to set the weight or branch length of an edge.
- **Show Weight** Toggle to display edge weights or branch lengths.
- **Measure Weights** Button to set edge weights based on coordinates.
- **Support** Text field to set confidence or support values (e.g. bootstrap).
- **Show Support** Toggle to display confidence or support values.
- **Probability** Text field to set probabilities for reticulate edges.
- **Show Probability** Toggle to display probabilities on reticulate edges.

#### 7.6 Edge Label Style Panel

This panel provides options for customizing the style and appearance of edge labels:

- **Font** Dropdown menu to select the font for edge labels.
- **Revert Font** Button to reset font settings to default.
- **Size** Combo box to specify or edit the font size for edge labels.
- **Revert Size** Button to reset size settings to default.
- **Bold** (B) Toggle bold formatting for edge labels.
- **Italic** (I) Toggle italic formatting for edge labels.
- **Underline** (U) Toggle underlining for edge labels.
- **Color** Color picker to set the text color of edge labels.
- **Revert Color** Button to reset text color to default.
- **Fill** Color picker to set the background fill color of edge labels.
- **Revert Fill** Button to reset the background fill color to default.

#### 7.7 Draw Phylogeny

This panel provides options for algorithmic drawing of the phylogeny.

There are three choices for layout:

- Rectangular,
- Circular,
- Radial.

There are three choices for scaling:

- To-scale phylogram,
- Early-branching cladogram,
- Late-branching cladogram.

The *Run* button re-runs the layout using the current settings.

#### 7.8 Layout Phylogeny

This panel provides tools for geometric transformations of the displayed phylogenetic tree or network:

- **Rotate Left** Rotate the entire phylogeny $90^\circ$ counterclockwise.
- **Rotate Right** Rotate the entire phylogeny $90^\circ$ clockwise.
- **Horiz. Flip** Reflect the phylogeny across a vertical axis.
- **Vert. Flip** Reflect the phylogeny across a horizontal axis.
- **Resize Mode** Toggle an interactive mode that allows resizing the layout by direct manipulation.
- **Layout Labels** Reposition node and edge labels to improve readability after transformations.

#### 7.9 Modify Phylogeny

This panel provides options for modifying the tree or network:

- **Magic Wand** Apply the most pertinent modification, based on the current selection of nodes and edges.
- **Induce** Keep only the part of the phylogeny induced by the current selection.
- **Declare Root** Assign a specific node as the root of the phylogenetic network, redirecting edges where necessary.
- **Merge Nodes** Combine multiple selected nodes into a single node while preserving connectivity.

- **Del. Thru Nodes** Delete nodes of indegree one and outdegree one.
- **Reverse Edges** Invert the direction of selected edges while maintaining network integrity.
- **Cross Edges** For any selected node of degree four, replace the node with two crossing edges.
- **Acceptor Edge** Declare an edge as the recipient of a horizontal gene transfer event. There can be at most one such edge per reticulation node.

### 8 Working with Nodes and Edges

#### 8.1 Creating Nodes and Edges

- In Edit Mode, drag along the canvas to create edges.
- In Edit Mode, shift-drag on selected nodes or edges to move them.
- Double-click (or long-press, on touch devices) to create new nodes.

#### 8.2 Transforming Nodes and Edges

- In Transform Mode, drag nodes to move them.
- Drag on edges to adjust their shape and control points.

### 9 Menus (Non-Mobile App Only)

#### 9.1 File Menu

The *File* menu contains the usual file-related items:

- **New...** Create a new document.
- **Open...** Open an existing document (file suffix `.psketch`). Additionally, you can import a tree or network from a file in Newick format (file suffix `.new`, `.tre` or similar) or from a file created by PhyloSketch1 (file suffix `.nexus`).
- **Recent** Access a list of recently opened files.
- **Export** Open the export submenu for saving data in different formats.
- **Image...** Export the current canvas as an image file.
- **Newick...** Export the current tree or network in Newick format.
- **Save...** Save the current document.
- **Page Setup...** Configure page layout and settings for printing.

- **Print...** Print the current document.
- **Close** Close the currently open document or window.
- **Quit** Exit the application.

#### 9.2 Edit Menu

The *Edit* menu items are:

- **Undo** Revert the last action.
- **Redo** Repeat the last undone action.
- **Cut** Remove the selected items and copy them to the clipboard.
- **Copy** Copy the selected items to the clipboard.
- **Copy Image** Copy an image of the current canvas to the clipboard.
- **Paste** Insert the contents of the clipboard into the current document.
- **Delete** Remove the selected nodes and/or edges.
- **Clear** Delete all nodes and edges from the canvas.
- **Apply Modification** Apply the most pertinent modification (among the items listed below), given the current selection of nodes and edges.
- **Remove Thru Nodes** Replace "thru nodes" (nodes with indegree 1 and outdegree 1) by direct edges.
- **Declare Root** Change the root of the tree or network by selecting a new node or edge.
- **Declare Acceptor Edge** For any reticulate node, exactly one of the incoming edges may be declared the transfer acceptor edge. If the selected edge is already declared an acceptor edge, it loses this property.
- **Mode** Switch between editing modes:
  - Edit Mode Enable adding and modifying nodes and edges.
  - Move Mode Allow repositioning of nodes and reshaping of edges.
- **Find...** Search for a node or edge by label.
- **Find Again** Repeat the previous search to locate the next matching node.
- **Add LSA Edges** Add edges representing Lowest Stable Ancestor (LSA) relationships between selected nodes.

#### 9.3 Layout Menu

The *Layout* menu provides items for customizing the layout and appearance of phylogenetic trees and networks. The following items are available:

- **Outlines** Toggle display of the tree or network as an outline.
- **Rotate Left** Rotate the tree or network 90 degrees to the left.
- **Rotate Right** Rotate the tree or network 90 degrees to the right.
- **Flip Horizontal** Flip the tree or network horizontally.
- **Flip Vertical** Flip the tree or network vertically.
- **Resize Mode** Enable or disable resize mode, allowing the layout to be resized and repositioned.
- **Layout Labels** Reset the layout of node and edge labels for better readability.
- **Layout Phylogeny** submenu:
  - **Apply** Lay out the phylogeny using the current settings.
  - **Radial Layout** Use a radial layout.
  - **Rectangular Layout** Use a rectangular layout.
  - **Circular Layout** Use a circular layout.
  - **To-Scale Phylogram** Lay out as a phylogram.
  - **Early-Branching Cladogram** Lay out as an early-branching cladogram.
  - **Late-Branching Cladogram** Lay out as a late-branching cladogram.

#### 9.4 View Menu

The *View* menu provides items for adjusting the appearance, scaling, and layout of the canvas:

- **Use Dark Theme** Toggle between light and dark themes for the application interface.
- **Increase Font Size** Increase the font size of labels and text in the canvas.
- **Decrease Font Size** Decrease the font size of labels and text in the canvas.
- **Zoom In** Zoom in on the canvas for a closer view.
- **Zoom Out** Zoom out of the canvas for a broader view.
- **Zoom To Fit** Adjust the zoom level to fit the entire phylogeny within the window.
- **Enter Full Screen** Switch the application to full-screen mode for an immersive view.

#### 9.5 Window Menu

The *Window* menu items are:

- **Set Window Size. . .** Set the exact size of the window.
- One menu item for each currently open document window.

#### 9.6 Help Menu

The *Help* menu contains the following items:

- **Check for Updates. . .** Check whether an update for the application is available.
- **About. . .** Show an information window about the program.
- **Help Window. . .** Show a window containing this help document.

### 10 Combining vs Transfer View

The reticulations in a rooted phylogenetic network can be drawn in two different ways.

- **Combining view** All incoming edges of a reticulation node are drawn as special edges that come together at the reticulation, representing similar amounts of incoming genetic material, as in the case of speciation-by-hybridization or reassortment.
- **Transfer view** One incoming edge has been declared a “transfer acceptor edge” and all other incoming edges are drawn as special edges that represent small amounts of incoming genetic material, as in the case of horizontal gene transfer.

By default, reticulations are drawn in a combining view. To obtain a transfer view, select one of the incoming edges and use the `Declare Acceptor Edge` menu item or button to declare it to be the transfer acceptor edge. If providing the network in extended Newick format, use ##H$i$ instead of #H$i$ to indicate which of the incoming edges of the $i$-th reticulation node is the acceptor edge.

This example shows the (complicated) published drawing of a network (Lescroart et al. [2023], Fig. S12E) and a (simpler) transfer view obtained using the *rectangular cladogram algorithm*:

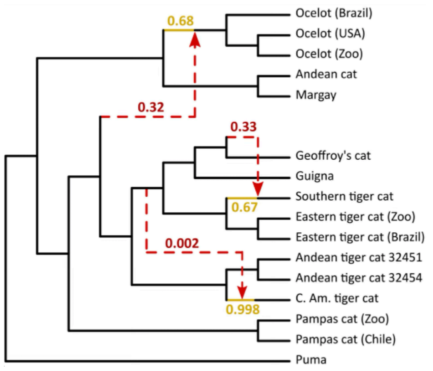

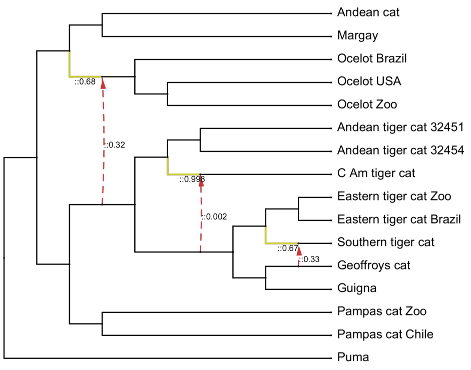

Here we show a captured hybridization network (Barley et al. [2022], Fig. 2) and a combining view obtained using the *rectangular cladogram algorithm*:

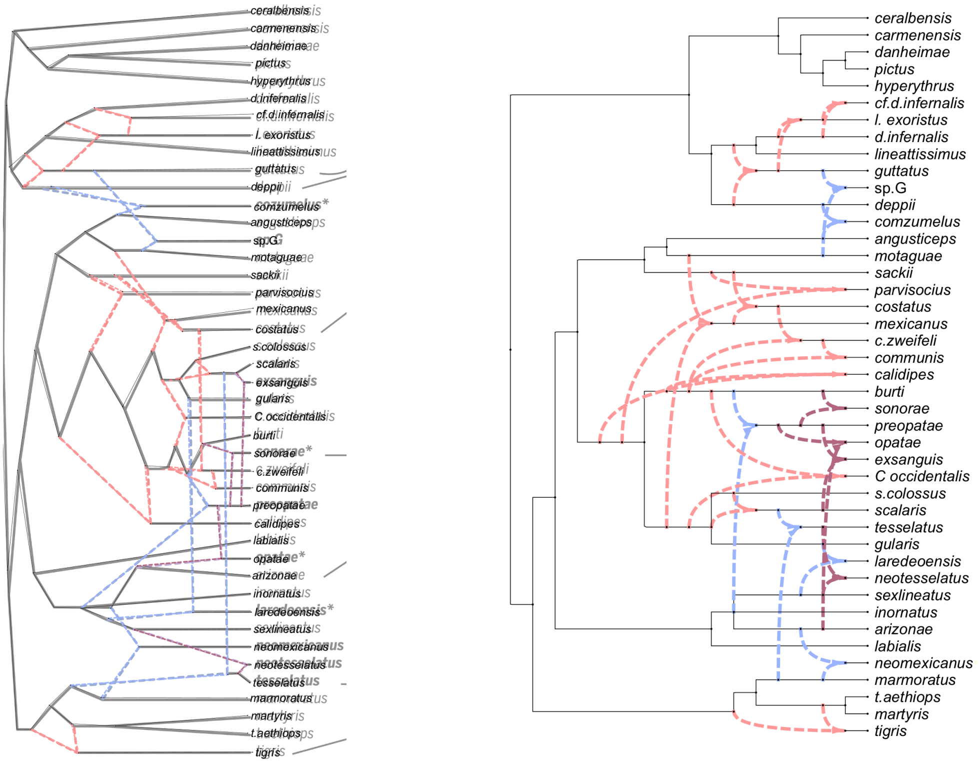

### 11 Example files

There are several example of image files and PhyloSketch files available online here:

`https://github.com/husonlab/phylosketch2/tree/main/examples`

### 12 Advanced features

The layout algorithm for rooted phylogenetic networks uses simulated annealing for nodes of large outdegree.but The default parameters are: start temperature = 1000, end temperature = 0.01, 1000 iterations per temperature step, cooling rate = 0.95.

We do not expose these parameters in the UI, however, if you really want to change these, then edit the properties file `PhyloSketch2.def` (its location is system specific, either `~/Library/Preferences/PhyloSketch2.def` or `~/.PhyloSketch2.def`) and write statements like this: `SA_DEFAULT_START_TEMPERATURE=2000`, `SA_DEFAULT_END_TEMPERATURE=1`, `SA_DEFAULT_ITERATIONS_PER_TEMPERATURE=100` and `SA_DEFAULT_COOLING_RATE=0.80` to change the values to 2000, 1, 100 and 0.8, say, respectively.

### 13 Support and Feedback

For issues, bug reports, or suggestions, please use the GitHub repository or the apps support page linked there.

#### 14 Third-Party Software

This software uses the Tesseract OCR engine, which is licensed under the Apache License, Version 2.0.
